# Breaking the bottleneck: semi-automated quantification of actin filament dynamics

**DOI:** 10.64898/2026.11.05.764262

**Authors:** Alexander D. Nobles, David M. Rutkowski, Cody R. Belden, Laura M. Haney, Amanda E. Davenport, Dimitrios Vavylonis, Jessica L. Henty-Ridilla

## Abstract

Actin reconstitution assays visualized by total internal reflection fluorescence (TIRF) microscopy have been crucial for defining mechanisms of filament assembly, disassembly, and higher-order organization. Yet quantitative analysis remains slow, labor-intensive, and subject to investigator bias. We developed SMAUG (<u>S</u>ingle filament <u>M</u>easurement and <u>A</u>nalysis <u>U</u>sing a <u>G</u>UI), a semi-automated image analysis interface that extends TSOAX filament segmentation and tracking with user-verified trace analysis of time-lapse TIRF images. SMAUG generates kymographs and quantifies filament nucleation, elongation, pausing, junctions, and filament-associated fluorescence while retaining measurements at the individual filament level. We tested the approach across actin reconstitution assays producing concentration-dependent polymerization, formin-stimulated elongation, capping-induced arrest, Arp2/3 branching, and fascin-mediated bundling. SMAUG reproduced manual measurement while reducing analysis time and revealed where dense networks and bundled filaments require user validation. Together, TSOAX and SMAUG provide a semi-automated approach for quantitative analysis of actin filament dynamics.

## Introduction

Actin filaments power many essential cellular processes, yet quantitative analysis of their dynamics remains slow, labor-intensive, and prone to measurement bias. Total internal reflection fluorescence (TIRF) microscopy paired with *in vitro* biomimetic reconstitution assays enables direct observation of filament assembly, disassembly, and higher-order structures^1–6^. However, extracting quantitative measurements from these assays remains a major bottleneck. Manual tracking of individual filament length and shape is time-consuming and limits the number of filaments that can be analyzed. Interactive tools such as JFilament^7^ facilitate this process by sequentially fitting deformable “active” contours along fluorescently labeled filaments but still require significant user involvement and remain susceptible to user selection bias.

Fully automated approaches offer an alternative but face the challenge of tracking individual filaments within evolving networks, where identifying the same filament across successive images can become ambiguous^8^. Recent advances include active-contour-based tracking of biopolymer networks in 2D and 3D^9^, analysis of microtubule growth and shrinkage with MTrack^10^, graph-based approaches for resolving self-intersections with KnotResolver^11^ and GraFT^12^, and machine-learning approaches that combine learned filament detection with temporal association^13,14^. Although these approaches provide powerful tools for automated filament analysis, efficient verification of individual filament traces and assessment of analysis parameters remain important challenges, particularly for 2D TIRF actin assays across different experimental conditions.

Here we demonstrate that efficient and systematic analysis of individual filament histories can be achieved by combining **T**rackable **S**tretching **O**pen **A**ctive **C**ontours (TSOAX)^9^ with a new user-friendly analysis interface, SMAUG (**S**ingle filament **M**easurement and **A**nalysis **U**sing a **G**UI). In this workflow, TSOAX identifies filaments in time-lapse images and links them across successive frames, while SMAUG enables users to inspect and verify the resulting filament traces, visualize filament behavior, generate kymo-graphs, and quantify filament nucleation, elongation, pausing, branching, and fluorescence intensity. We tested this approach across diverse reconstituted actin assays that produce distinct filament behaviors and dimensions, including concentration-dependent polymerization, formin-stimulated elongation, capping-induced arrest, Arp2/3-mediated branching, and fascin-mediated bundling.

Across these assays, SMAUG-assisted analysis reproduced measurements obtained by manual analysis while substantially reducing analysis time and retaining measurements at the individual filament level. Testing increasingly complex filament behaviors also revealed where automated segmentation benefits from user verification, including dense branched networks and highly bundled filaments. Together, TSOAX and SMAUG provide a semi-automated approach for quantitative analysis of actin filament dynamics that reduces manual selection and measurement while retaining user oversight where image interpretation remains essential. This approach can be applied to different actin regulators and experimental conditions and provides a way to quantify individual filament behaviors across reconstituted assays.

## Results

### TSOAX and SMAUG enable semi-automated analysis of actin filament dynamics

We first established a semi-automated approach for quantifying actin filament dynamics from time-lapse microscopy images of 1 µM actin actively polymerizing (Figure 1). In conventional analysis, individual filaments are identified and traced across successive frames of a movie to measure filament number as a readout of nucleation, changes in filament length to calculate elongation rates, and other features of filament dynamics depending on the experimental question (Figure 1A). These measurements depend on manual selection of filaments and time points, and typically only a few frames are measured per filament, making the analysis time-consuming and limiting the number of filaments that can be reasonably analyzed. To streamline this process, we combined the SMAUG analysis interface developed here with TSOAX filament segmentation and tracking^9^ (Figure 1B, C).

**Figure 1.**
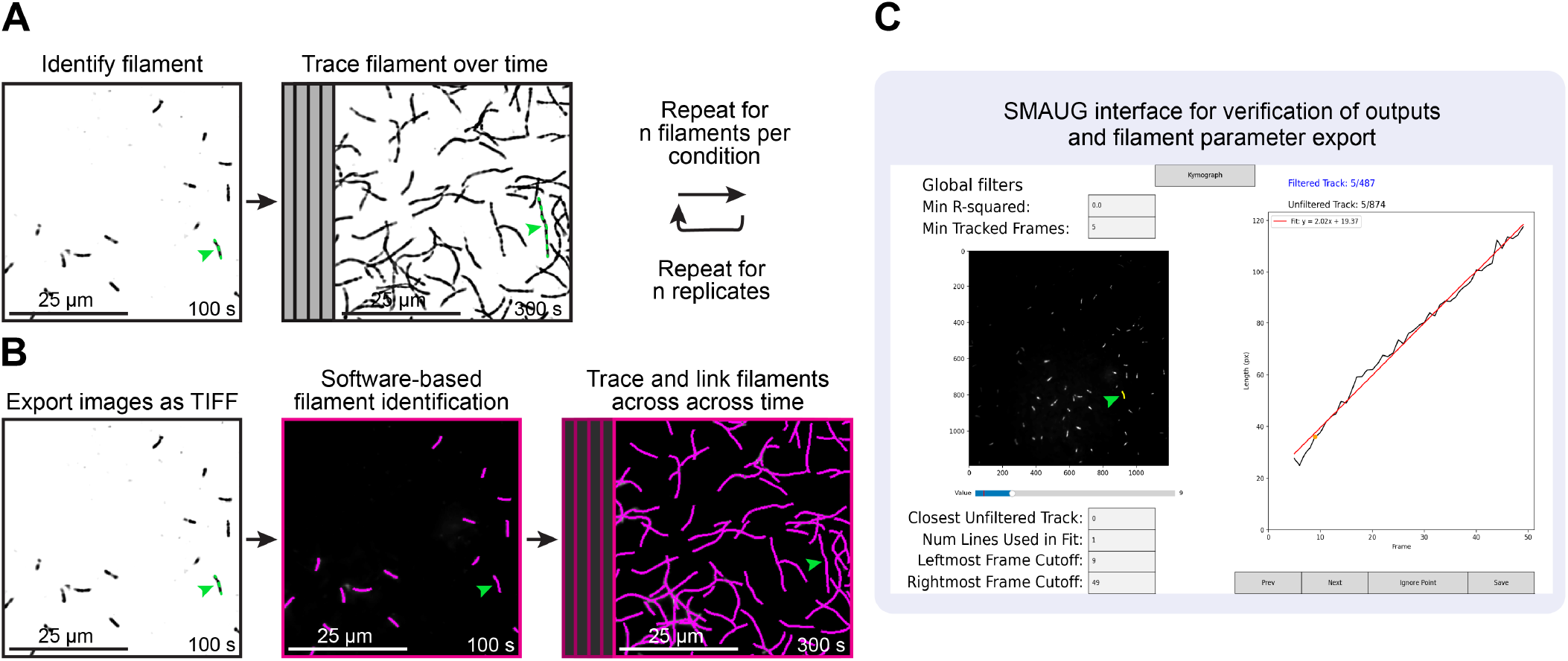
Comparison of manual and SMAUG-assisted image analysis of actin filaments. **(A)** Conventional manual analysis of actin filament elongation from TIRF microscopy images. Individual filaments are identified and manually traced across successive time points to measure changes in filament length. This process is repeated for multiple filaments per condition and across experimental replicates. The green arrow identifies an example filament followed throughout the analysis. **(B)** TSOAX segmentation and SMAUG-assisted analysis of the same TIRF images presented in (A). TIRF images are exported as TIFF files and analyzed using TSOAX to identify filament segments (magenta), which SMAUG displays for user validation. **(C)** SMAUG graphical user interface (GUI) is used to verify filament trace accuracy and export measurements. Example filament (green arrow) length is plotted over time, and the red line shows the linear fit used to calculate the mean filament elongation rate from its slope. Scale bars, 25 µm.

In this workflow, time-lapse images are exported as TIFF files and first analyzed with TSOAX to identify filaments and link them across successive frames. Initial filament segmentation is performed using the TSOAX initialization function, followed by refinement of the segmented filaments using the deform function before tracking across the movie. The resulting traces are exported from TSOAX and imported into SMAUG, where they are displayed for user inspection and verification. SMAUG reports the number of frames across which each filament was successfully tracked, allowing users to assess whether individual traces accurately follow the filament and remain reliable over the desired analysis period. Users can also manually exclude individual frames containing tracking errors while retaining the remaining measurements for that filament. This is particularly useful when a filament tip briefly moves out of focus, preventing accurate tracking in an individual frame without compromising measurements from the rest of the movie.

SMAUG also provides options for refining which filaments and time points are included in the final analysis. Users can filter traces based on a user-defined minimum number of successfully tracked frames and then verify that the selected traces are accurately tracked over the chosen period. Users can also exclude filaments that cross the edge of the field of view (FOV) when measurements require tracking the complete filament length, such as calculating elongation rates. However, these filaments may still be retained for measurements of filament number or nucleation, for which the presence of a filament can be sufficient. In addition, users can define the time window used for analysis to focus on periods when filament behavior can be reliably measured, such as after sufficient actin has accumulated or following the addition of a regulatory protein. For measurements of filament elongation, SMAUG allows users to fit up to 6 independent linear segments to different portions of an individual filament’s length versus time record, enabling changes in growth rate to be quantified within a single filament. Following trace verification and selection of analysis parameters, measurements including filament elongation rates and other quantitative features can be exported for further analysis.

### Semi-automated filament tracing reproduces manual measurements of actin assembly

We first asked whether TSOAX-based image segmentation followed by SMAUG analysis could reproduce measurements obtained by manual examination of actin polymerization from time-lapse TIRF microscopy assays. In standard 1 µM actin polymerization reactions, TSOAX identified and linked individual filaments across successive frames, allowing the same filaments to be followed over time (Figure 2A, B). Actin filament nucleation, measured as the filament count at 150 s, was similar between manual analysis and SMAUG analysis, with 102.4 ± 57.5 (SD) filaments per FOV measured manually compared with 102.7 ± 56.9 filaments per FOV measured by SMAUG (p = 0.99; Figure 2C). Thus, filament counts from the TSOAX and SMAUG analysis were comparable to those obtained by the direct inspection of the images.

**Figure 2.**
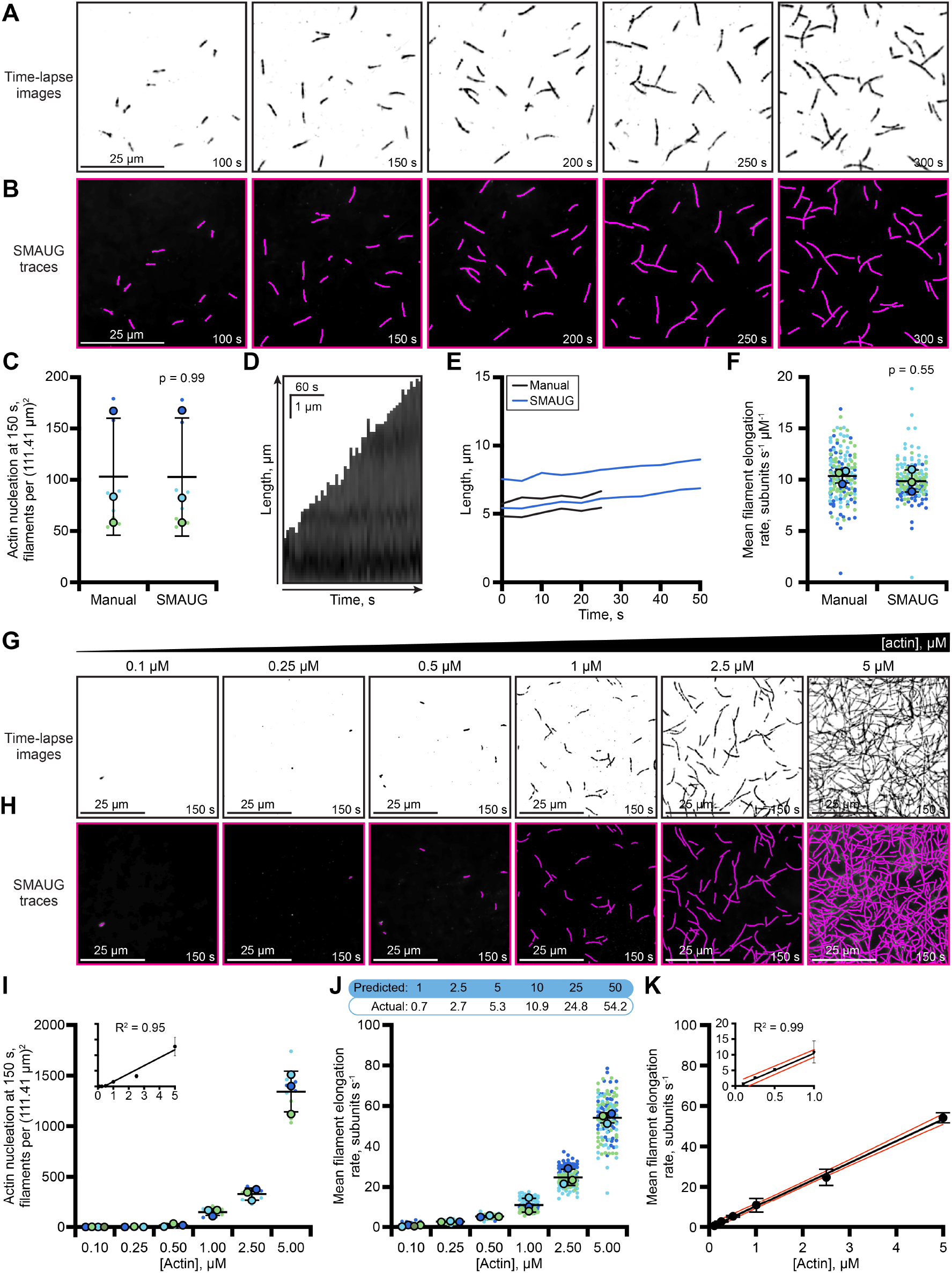
Semi-automated filament tracing reproduces manual measurements of actin assembly. **(A)** Representative time-lapse images of TIRF reactions containing 1 µM actin (10% AlexaFluor-488, 3.6 nM biotin-actin). **(B)** TSOAX-generated filament traces (magenta) linked across successive frames for analysis in SMAUG. **(C)** Number of filaments per field of view (FOV) at 150 s measured manually and by SMAUG. For each condition, all filaments present in each FOV were counted across N = 3 independent reactions, with n = 9 FOVs total. Small dots represent individual FOVs and large dots indicate reaction means. Colors denote independent reactions. **(D)** SMAUG-generated kymograph of an individual actin filament from (A). **(E)** Representative filament length versus time plots generated from manual (black) and SMAUG (blue) measurements. **(F)** Mean filament elongation rates measured manually or by SMAUG. 17 filaments per FOV, were measured across N = 3 independent reactions (n = 153 filaments). Small dots represent individual filaments and large dots indicate reaction means. Colors denote independent reactions. **(G)** Representative time-lapse TIRF images of reactions containing increasing concentrations of actin (10% AlexaFluor-488, 3.6 nM biotin-actin). **(H)** TSOAX-generated filament traces (magenta), used for analysis in SMAUG. **(I)** Filament nucleation across actin concentration gradient. Each dot represents the filament count per FOV (n = 9, except 0.1 µM actin, where n = 12) from N = 3 independent experiments, except 0.1 µM, where N = 4. Inset, linear regression of mean nucleation values versus concentration (R^2^ = 0.95). **(J)** Mean filament elongation rates at varied actin concentrations. 17 filaments per FOV, or all filaments if fewer were measured across N = 3 independent reactions (n = 153 filaments), except 0.1 µM actin (n = 70 filaments total) and 5 µM actin (n = 148 filaments total). 1 µM actin included N = 3 independent reactions. Small dots represent individual filaments and large dots indicate reaction means. Colors denote independent reactions. **(K)** Linear regression of mean filament elongation rates from (J) (R^2^ = 0.99). Red lines indicate the 95% confidence interval of the regression line. Inset, expanded view of measurements at actin concentrations below 1 µM, showing that these values fall within the confidence interval of the regression line. Statistical comparisons between manual and SMAUG measurements were performed using Welch’s t-test (2C, p = 0.99; 2F, p = 0.55).

We compared mean filament elongation rates from the two approaches. Because the TSOAX traces preserve filament position across successive frames, SMAUG could generate kymographs from individual tracked filaments (Figure 2D). In these kymographs, filament growth is represented by the slope of the filament trace over time, providing a visual representation of the same growth behavior quantified from filament length versus time. Representative length versus time plots showed similar slopes for manual and SMAUG measurements (Figure 2E). Across 153 filaments, the mean elongation rate was 10.4 ± 0.67 subunits s^-1^ µM^-1^ measured manually compared with 9.9 ± 1.1 measured by SMAUG (p = 0.55; Figure 2F). SMAUG measurements incorporated more time points (frames) per filament than were typically measured for manual measurements, often >10 frames compared with 5-7 for manual measurements (Supplementary Figure 1A), providing more consistent sampling of filament growth without altering the measured elongation rates.

We next asked whether the same analysis could be applied across a range of actin concentrations spanning from 0.1 to 5 µM (Figure 2G, H). Filament nucleation increased linearly with actin concentration (R^2^ = 0.95; Figure 2I), as did mean elongation rates (R^2^ = 0.99; Figure 2J, K), consistent with the expected concentration dependence of actin polymerization^8–11^. Notably, SMAUG-derived measurements of mean elongation rate, filament dissociation rate (0.35 subunits s^-1^), and the critical concentration for actin assembly (0.03 µM) were consistent with the 95% confidence intervals of regression fits, and comparable to previously reported measurements (1.4 ± 0.8 subunits s^-1 15^, and 0.12 ± 0.07 µM^15^, respectively; Figure 2J). At lower actin concentrations, the number of measurable filaments was limited by filament abundance within the FOV. At lower actin concentrations, the number of measurable filaments was limited by filament abundance within the FOV. In contrast, at 5 µM actin, rapid growth and increased filament density made individual traces more difficult to resolve, although elongation rates could still be measured. Thus, TSOAX and SMAUG reproduced the expected concentration dependence of both filament nucleation and elongation across the range tested, while the highest concentration may begin to challenge filament segmentation and tracking.

### SMAUG analysis of formin-stimulated changes in actin assembly

We next tested whether SMAUG could resolve changes in actin filament behavior produced by actin regulatory proteins. Profilin 1 (PFN1) suppresses spontaneous actin nucleation^19–21^. In contrast, the formin mDia1 promotes filament formation and, in the presence of PFN1, supports rapid processive actin filament elongation^20,22-27^. Therefore, we compared TIRF reactions containing 1 µM actin alone, actin with 1 µM PFN1, actin with 5 nM mDia1(FH1-C), and actin containing both PFN1 and mDia1(FH1-C) (Figure 3A, B). PFN1 reduced the number of filaments detected at 150 s from 58.6 ± 15.7 in actin alone reactions to 27.2 ± 18.0 filaments per FOV (Figure 3C; p = 0.99). SMAUG generated kymographs (Figure 3D) and representative length versus time plots (Figure 3E) that captured differences in filament growth across these conditions. Mean elongation rates were lower in the presence of PFN1, decreasing from 9.8 ± 0.4 to 6.4 ± 0.8 subunits s^-1^ µM^-1^ (p = 0.96; Figure 3F). The addition of mDia1(FH1-C) substantially increased the number of filaments detected to 690.1 ± 398.7 filaments per FOV (p = 0.05; Figure 3C), while mean elongation rates remained similar to actin alone reactions at 10.4 ± 1.0 subunits s^-1^ µM^-1^ (p = 0.99; Figure 3D-F). In contrast, combining PFN1 with mDia1(FH1-C) elevated the number of actin filaments to 271.1 ± 261.5 filaments per FOV (p = 0.29; Figure 3C) and significantly increased the mean elongation rate to 41.2 ± 16.4 subunits s^-1^ µM^-1^ (p < 0.01; Figure 3D-F), within the reported range for PFN1 and mDia1-stimulated actin assembly^20,23,28-31^.

**Figure 3.**
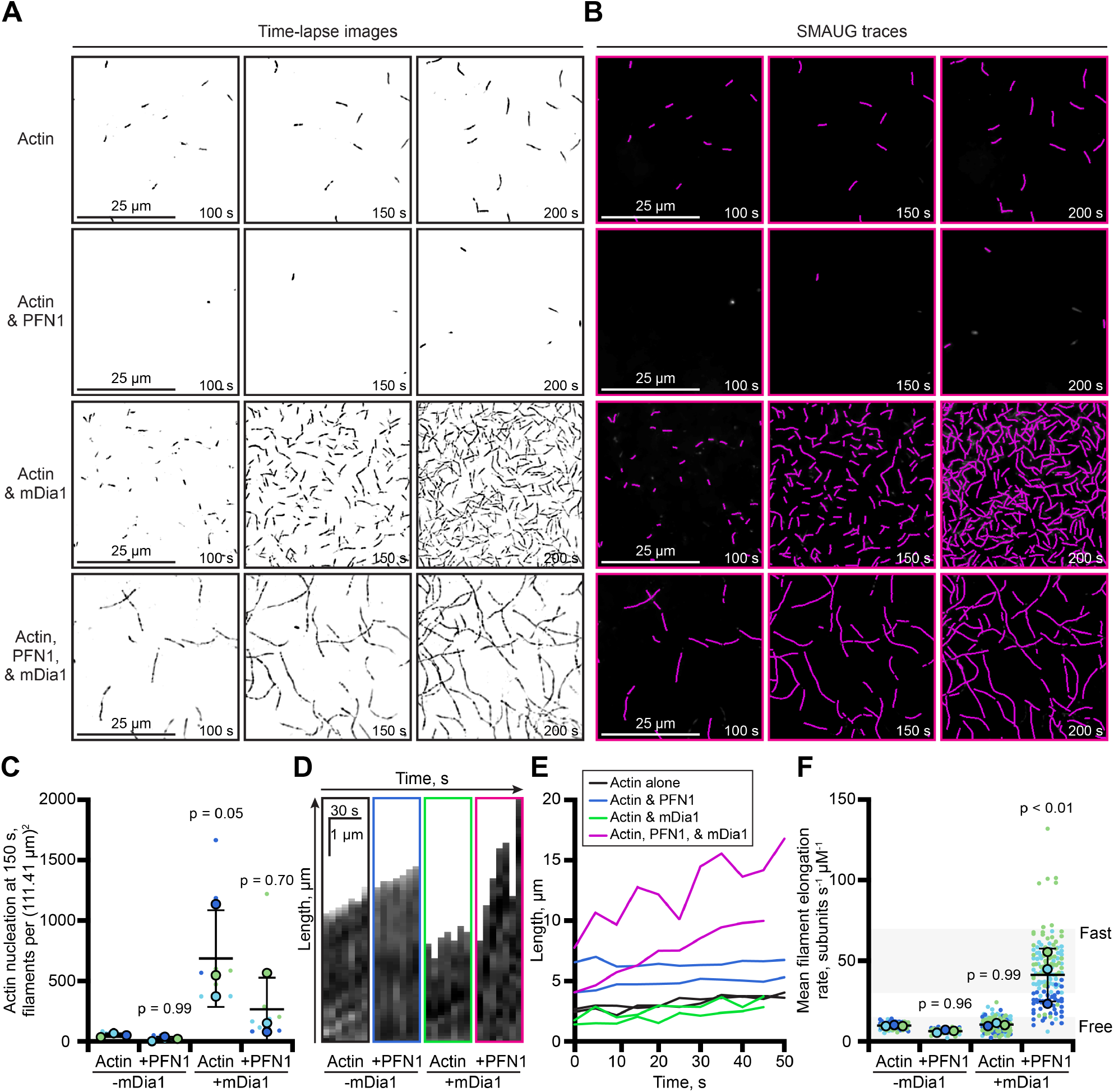
SMAUG analysis captures formin-dependent increases in actin nucleation and elongation. **(A)** Representative time-lapse TIRF images of 1 µM actin (10% AlexaFluor-488, 3.6 nM biotin-actin) polymerizing in the absence or presence of 1 µM profilin-1 (PFN1) or 5 nM mDia1(FH1-C), or the combination of both proteins for fast elongation. **(B)** TSOAX-generated filament traces (magenta) for the images in (A). **(C)** Number of filaments per field of view (FOV) at 150 s. All filaments were counted across N = 3 independent reactions (n = 9 FOVs). Small dots represent individual FOVs and large dots indicate reaction means. Colors denote independent reactions. **(D)** SMAUG-generated kymographs for the indicated conditions. **(E)** Representative filament length versus time plots. **(F)** Mean filament elongation rates. 17 filaments per FOV were measured across N = 3 independent reactions (n = 153; except actin & PFN1, n = 104; and actin, PFN1, & mDia1, n = 146). Small dots represent individual filaments and large dots indicate reaction means. Colors denote independent reactions. Statistical analysis was performed by one-way ANOVA with Tukey’s post-hoc test, with comparison to actin alone for reactions containing PFN1, mDia1, and both PFN1 and mDia1 (3C, p = 0.99, p = 0.05, and p = 0.70; 3F, p = 0.96, p = 0.99, and p < 0.01, respectively). Data are mean ± SD. All scale bars, 25 µm.

Notably, individual filament measurements revealed heterogeneity in elongation rates within the same reaction, particularly in the PFN1-mDia1(FH1-C) condition, where filaments displayed a broad range of rates (Figure 3F). Because elongation rates differ between formin-associated and formin-free filaments, these measurements may provide a means of identifying differences in formin occupancy within a population of filaments^20,28,32,33^. Thus, SMAUG preserves the measurements at the individual filament level while allowing these distributions to be quantified across many filaments. However, at the high signal density produced by mDia1, increased signal overlap introduced challenges for automated segmentation and tracking.

### SMAUG analysis resolves individual capping protein-induced pauses in actin filament growth

We next tested whether SMAUG could resolve filament capping events and quantify the resulting pauses in actin assembly (Figure 4A, B). The addition of 10 nM capping protein (CP) reduced the number of filaments detected at 150 s from 301.6 ± 155.0 to 95.1 ± 28.2 filaments per FOV (p = 0.14; Figure 4C). Kymographs and filament length over time plots revealed distinct transitions from active elongation to periods of little or no net growth in the presence of CP (Figure 4D-F; red arrows), consistent with previous studies^34-38^. Across all tracked filaments, mean elongation rates were reduced from 9.6 ± 1.3 to 5.6 ± 0.9 subunits s^-1^ µM^-1^ for reactions including CP (p = 0.02; Figure 4G). Importantly, the analysis also captured rare uncapping events where filaments resumed elongation after a period of arrest (Figure 4F; green arrow). Elongation rates during active growth were similar before capping and after growth resumed, indicating that the same filament could be followed through active, paused, and resumed growth states. Inspection of individual traces in SMAUG allowed users to verify that these transitions reflected changes in filament growth rather than tracking errors, such as those that can occur when a growing filament end encounters a junction. This verification also provides a way to assess whether the imaging interval is appropriate for resolving these transitions, since intervals that are too long or too short may limit the ability to distinguish changes in growth from tracking errors.

**Figure 4.**
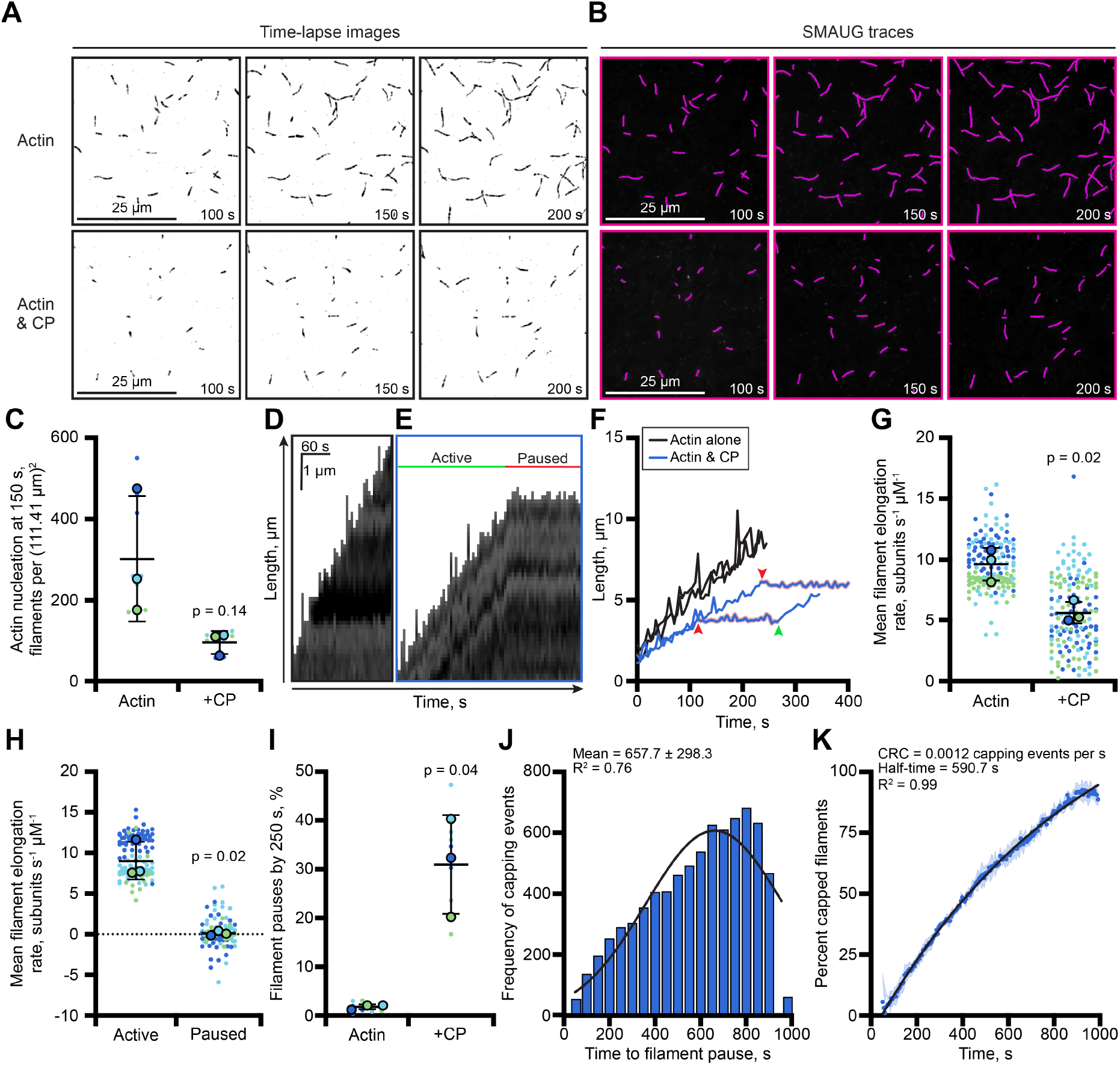
SMAUG analysis resolves individual capping protein-induced pauses in actin filament growth. **(A)** Representative time-lapse TIRF images of 1 µM actin (10% AlexaFluor-488, 3.6 nM biotin-actin) polymerizing in the absence or presence of 10 nM capping protein (CP). **(B)** TSOAX-generated filament traces (magenta) for the images in (A). **(C)** Filament nucleation per field of view (FOV) at 150 s. All filaments were counted across N = 3 independent reactions (n = 9 FOVs). Small dots represent individual FOVs and large dots indicate reaction means. Colors denote independent reactions. **(D)** SMAUG-generated kymograph of an actin filament from a control reaction containing actin alone. **(E)** SMAUG-generated kymograph of an actin filament from a reaction containing CP. Active and paused filament elongation are labeled. **(F)** Representative filament length versus time plots. Red arrows indicate the onset of capping events, and the green arrow indicates a rare uncapping event where filament elongation resumed. **(G)** Mean filament elongation rates. 17 filaments per FOV, or all filaments if fewer, were measured across N = 3 independent reactions (n = 153 for actin alone; n = 150 for actin & CP). Small dots represent individual filaments and large dots indicate reaction means. Colors denote independent reactions. **(H)** Mean filament elongation rates during active and paused growth. Small dots represent individual filaments and large dots indicate reaction means. Colors denote independent reactions (N = 3; n = 115 filaments). **(I)** Percent of filaments with paused growth by 250 s. Each small dot represents one FOV (n = 9 FOVs from N = 3 independent reactions); large dots indicate reaction means and colors denote independent reactions. **(J)** Distribution of time to CP-induced pause events. Gaussian fit; R^2^ = 0.76; mean ± SD. **(K)** Percentage of capped filaments over time. Data were averaged within each independent reaction before fitting to determine the capping rate constant (CRC = 0.0012 capping events s^-1^) and corresponding half-life (590.7 s). One representative fit is shown (R^2^ = 0.99); data represent n = 9 FOVs from N = 3 independent reactions (mean ± SD). All scale bars, 25 µm. Statistical analysis, Welch’s t-test with actin alone control (4C, p = 0.14; 4G, p = 0.02; 4H, p = 0.02; 4I, p = 0.04).

Individual filament measurements provide a more direct view of these capping events. SMAUG identified transitions from active elongation to paused growth and allowed elongation rates to be measured separately during each state. Active filaments in CP-containing reactions elongated at 9.0 ± 2.3 subunits s^-1^ µM^-1^ compared with 0.1 ± 0.3 subunits s^-1^ µM^-1^ during paused periods (p = 0.02; Figure 4H). By 250 s, 31.0 ± 10.1 percent of filaments in CP-containing reactions had entered a paused state compared with 1.8 ± 0.5 percent in actin alone control reactions (p = 0.04; Figure 4I). The distribution of individual times until pausing captured the timing of CP-induced filament growth arrest (Figure 4J). We then quantified the accumulation of paused filaments over time and fit these data to determine the capping rate constant (0.0012 capping events per s), from which we calculated a capping half-life of 590.7 s (Figure 4K), consistent with previous estimations^34,35^. Together, these measurements show that individual filaments can be followed through active growth, arrest, and rare regrowth events, providing access to kinetic measurements that are difficult to capture systematically from endpoint measurements or manual analysis.

### Segmentation and analysis of Arp2/3 actin assembly and branched networks

We next tested TSOAX and SMAUG under conditions that pose a particularly difficult challenge for filament segmentation: Arp2/3 mediated branched actin networks containing frequent, dynamic filament-filament intersections (Figure 5A, B). Although TSOAX identifies filament intersections in these images (yellow dots), distinguishing *bona fide* branch points from overlapping filaments becomes challenging as network density and filament dynamics increase. These reactions provided a test of filament tracking in a complex network and define the limitations of automated branch detection with this analysis workflow.

**Figure 5.**
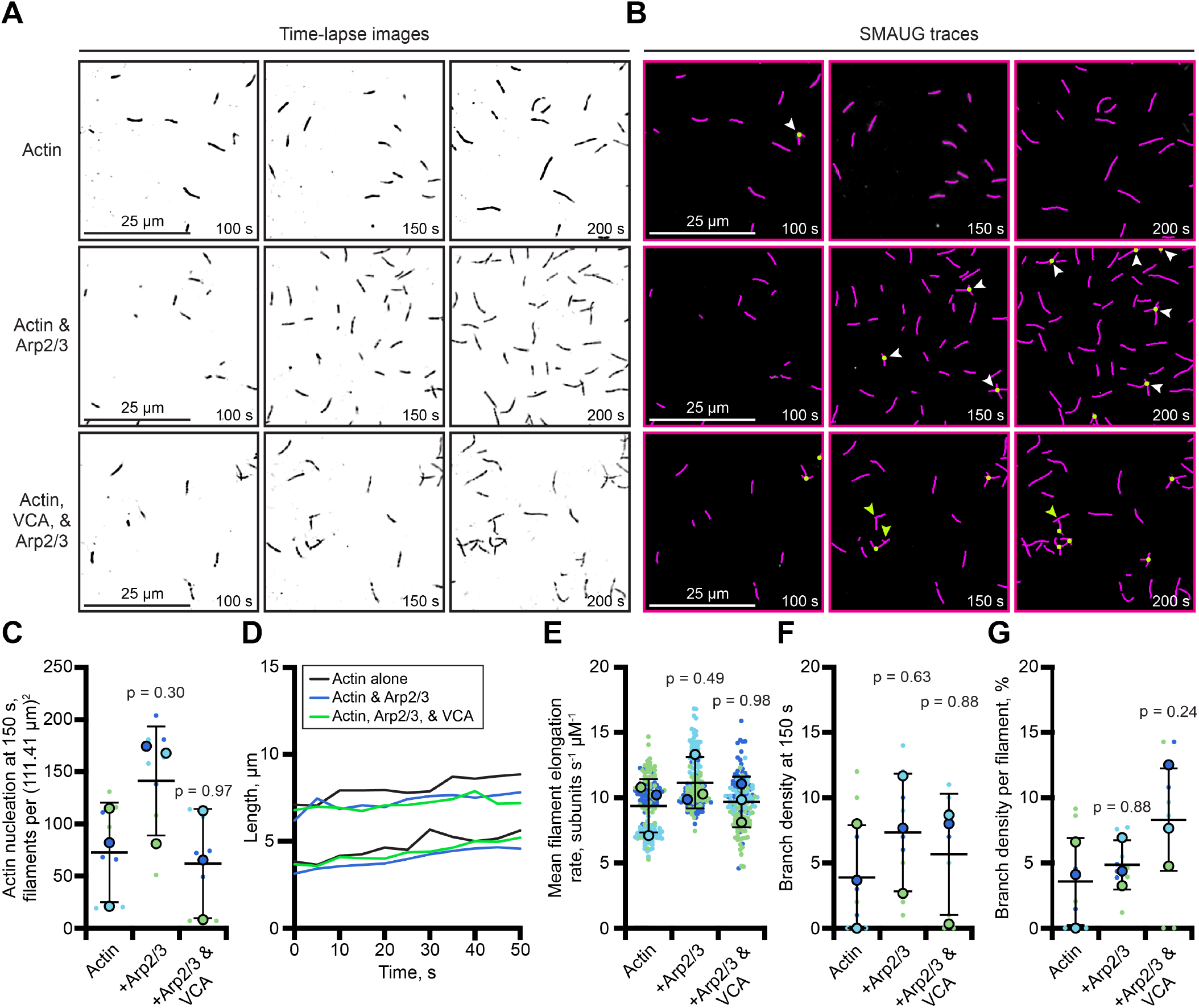
Segmentation and analysis of Arp2/3-dependent actin assembly and branching. **(A)** Representative time-lapse TIRF images of 1 µM actin (10% AlexaFluor-488, 3.6 nM biotin-actin) alone or supplemented with 100 nM Arp2/3 complex with and without 500 nM VCA (Arp2/3 complex activator). **(B)** TSOAX-generated filament traces for the images in (A). Yellow dots, filament-filament junction points identified by TSOAX. Yellow arrows, missed branch points. White arrows, misidentified branch points. **(C)** Filament nucleation per field of view (FOV) at 150 s. All filaments were counted across N = 3 independent reactions (n = 9 FOVs). Small dots represent individual FOVs and large dots indicate reaction means. Colors denote independent reactions. **(D)** Representative filament length versus time plots. **(E)** Mean filament elongation rates. 17 filaments per FOV were measured across N = 3 independent reactions (n = 153 filaments). Small dots represent individual filaments and large dots indicate reaction means. Colors denote independent reactions. **(F)** Number of filament-filament junctions per FOV at 150 s. All filament-filament crossing events were counted across N = independent reactions (n = 9 FOVs). Dots colored and represented as in (C). **(G)** Filament-filament junctions normalized to the number of filaments per FOV at 150 s. All filament-filament crossing events were counted across N = 3 independent reactions (n = 9 FOVs). Dots colored and represented as in (C). Statistical analysis was performed by one-way ANOVA with Tukey’s post-hoc test, with comparisons to actin alone with reactions containing Arp2/3 or Arp2/3 & VCA (5C, p = 0.30 and 0.97; 5E, p = 0.49 and p = 0.98; 5F, p = 0.63 and p = 0.88; 5G, p = 0.88 and p = 0.24, respectively). Data are mean ± SD. All scale bars, 25 µm.

We examined actin assembly in reactions containing 1 µM actin alone, actin with 100 nM Arp2/3, or actin with 100 nM Arp2/3 and 500 nM VCA to activate Arp2/3 dependent branching^39-44^. Filament number at 150 s was similar across conditions, with 72.3 ± 47.7 filaments per FOV in actin alone reactions, 140.9 ± 52.3 in reactions containing Arp2/3 alone (p = 0.3), and 61.8 ± 52.2 in reactions containing Arp2/3 and VCA (p = 0.97; Figure 5C). SMAUG generated filament length versus time plots and elongation measurements across all conditions (Figure 5D, E). Mean elongation rates were similar between actin alone (9.4 ± 2.0 subunits s^-1^ µM^-1^), Arp2/3 alone (11.2 ± 1.9 subunits s^-1^ µM^-1^; p = 0.49) and Arp2/3 with VCA (9.7 ± 1.5 subunits s^-1^ µM^-1^; p = 0.98; Figure 5E).

We next asked whether the identified filament junctions could be used to quantify branch density. TSOAX identified inter-sections where segmented filaments overlapped or connected, but inspection of the underlying images showed that some *bona fide* branch points were missed (Figure 5B; yellow arrows), while filament crossovers were also misidentified as intersections (Figure 5B; white arrows). Thus, the number of identified intersections did not directly report branch formation. The number of intersections per FOV increased from 3.9 ± 4.0 in actin alone to 7.3 ± 4.5 in Arp2/3 alone (p = 0.63) and 5.7 ± 4.6 with Arp2/3 and VCA (p = 0.88; Figure 5F). Normalizing these measurements to filament number did not resolve the issue: branch density increased from 3.6 ± 3.3 percent for actin alone to 4.9 ± 1.9 percent for Arp2/3 alone (p = 0.88) and 8.3 ± 3.9 percent for Arp2/3 with VCA (p = 0.24; Figure 5G). Although the analysis identified filament intersections, dense and dynamic networks could confound the assignment and linking of individual filaments, limiting the accuracy of these measurements as a direct readout of branching. Instead, branch density provides a useful starting point for user validation, reducing the amount of the FOV that must be inspected manually while retaining user judgement when filament identity and branch structure cannot be unambiguously resolved by segmentation alone.

### sIntensity-based analysis of actin filament bundles

We next asked whether SMAUG could quantify changes in filament fluorescence intensity under conditions that promote actin filament bundling. Addition of 1 µM fascin to 1 µM actin reactions produced visibly thicker and brighter filament structures compared with actin alone reactions (Figure 6A, B), consistent with fascin-mediated filament bundling^45–48^. SMAUG generated kymographs from the traced filaments, and although segmentation reduced bundled structures to a single pixel-wide “backbone” for tracking, intensity was extracted from the corresponding region of the original image, preserving the fluorescence signal within the tracked structure (Figure 6C, D). We therefore measured fluorescence intensity along each segmented filament to quantify changes in filament-associated fluorescence. At 600 s, mean filament intensity was 212.3 ± 22.3 AU in actin alone reactions and elevated to 305.6 ± 44.2 AU in reactions containing fascin (p = 0.05; Figure 6E). This measurement can also be followed over time, allowing changes in filament-associated fluorescence to be quantified throughout the reaction rather than only a defined time point. Filament intensity remained relatively stable over time across filaments in actin alone reactions, whereas intensity progressively increased for filaments polymerizing in the presence of fascin (Figure 6F). Thus, SMAUG can quantify changes in filament-associated fluorescence even as filaments overlap and form thicker structures.

**Figure 6.**
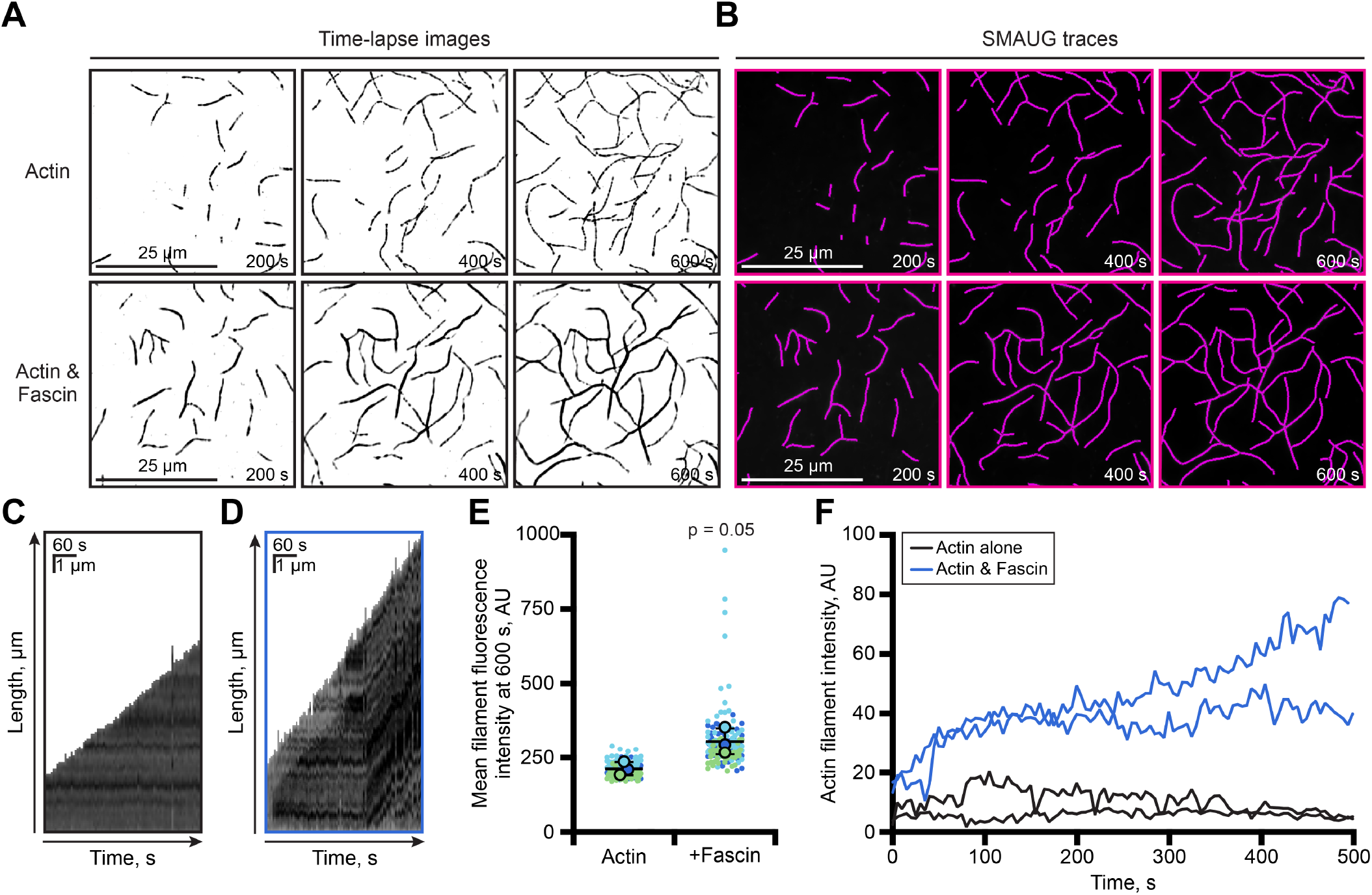
Measurement of actin filament bundling by fluorescence intensity. **(A)** Representative time-lapse TIRF images of 1 µM actin (10% AlexaFluor-488, 3.6 nM biotin-actin) polymerizing in the absence or presence of 1 µM fascin. **(B)** TSOAX-generated filament traces for the images in (A). **(C)** SMAUG-generated kymograph of an actin filament from a control reaction containing actin alone. **(D)** SMAUG-generated kymograph of an actin filament from a TIRF reaction containing fascin. Darker color on kymograph indicates increased fluorescence intensity. **(E)** Mean filament fluorescence intensity at 600 s. Intensity was measured along each segmented filament in the original image. 17 filaments per field of view (FOV) were measured across N = 3 independent reactions (n = 153 filaments). Small dots represent individual filaments and large dots indicate reaction means. Colors denote independent reactions. Statistical analysis was performed by Welch’s t-test compared to actin alone control (p = 0.05). Data are mean ± SD. **(F)** Mean filament fluorescence intensity over time. Lines represent the mean intensity of individual tracked filaments over time. Scale bars, 25 µm.

## Discussion

Quantitative image analysis has been a rate-limiting step for single-molecule assays of actin dynamics for more than two decades^3,49^. These reconstituted assays have been extremely useful for defining the biochemical mechanisms that control actin assembly, but manual filament selection and measurement is slow, inherently subjective, and can limit the experimental scale based on what can be reasonably analyzed. Here, we combine TSOAX filament segmentation with SMAUG analysis to make these measurements faster and more consistent while retaining user oversight. We found that the approach reproduced manual measurements of filament nucleation and elongation rates while substantially reducing the time required for analysis (Supplementary Figure 1B). This allowed us to analyze more filaments and fields of view without changing the basic measurements that are commonly made from these assays.

We tested the analysis under conditions that progressively challenged filament image segmentation. Formin-mediated assembly was one of the first places where we saw the limits of the approach. mDia1 rapidly increases the number of filaments in the field, and the resulting overlap and increased fluorescence signals made the TSOAX traces less consistent. Even under these conditions, SMAUG allowed us to measure substantially more rapidly growing filaments in less time than manual analysis, and the measured rates were within the range expected for these reactions^15,16,23,50^. These experiments also highlighted an important limitation of automated analysis: software cannot compensate for problems that originate with the imaging itself. At high polymer concentrations, or when filaments become very long, filaments can move into the imaging plane and appear to elongate. A user can recognize this by inspecting the images, but the software cannot distinguish this apparent elongation from true polymerization. Such events may therefore contribute to the small population of unusually fast measurements in the automated data. Thus, manual inspection of these reactions remains important, particularly when imaging conditions produce substantial filament overlap or movement out of the imaging plane.

The Arp2/3 reactions highlighted a related limitation of image segmentation: a junction identified in a 2D TIRF image is not necessarily a *bona fide* branch. Overlapping filaments can produce the same segmentation result as a branch point, causing the software to occasionally misidentify signal crossovers as branches. This may explain why the automated analysis reported more branching in the reactions containing Arp2/3 alone than in reactions containing Arp2/3 activated by VCA. The limitation was not in finding junctions, but in understanding what those points represented biologically. Thus, we recommend using junction detection to identify candidate branch points for rapid user verification, which is substantially faster than searching the entire imaging field manually. We observed a related limitation with fascin-bundled filaments. Although SMAUG provides a consistent location for measuring fluorescence intensity, the segmented backbone does not capture the full width of a bundle, and increasingly bundled filaments become difficult to distinguish as individual filaments. Thus, SMAUG can identify and quantify these structures, while their organization still requires user review.

Capping protein provided a particularly useful demonstration of the value of this approach. SMAUG identified individual filaments that were actively growing, detected when growth stopped, measured the duration of the pause, and captured relatively rare filament regrowth events. Because these events can be identified across many filaments, the data can be used to examine capping kinetics over time and (with appropriate experimental conditions) estimate association and dissociation parameters. This creates a direct link between microscopy-based reconstitution assays and more traditional biochemical measurements. Importantly, the approach also preserves information about the behavior of individual filaments that is lost when only the final filament population is measured. The ability to identify and quantify these heterogeneous single filament events across large numbers of filaments may therefore allow kinetic behaviors that are difficult to resolve by manual analysis to be examined more systematically.

A similar framework could also be extended to other measurements of cytoskeletal dynamics. Filament disassembly should be amenable to similar analysis, although severing produces many short fragments that become difficult to track individually. Under these conditions, filament number or total filament mass may provide more robust measurements than counting individual severing events^16,51-53^. The approach could also be applied to microtubule assays, although several features of these experiments would require additional analysis capabilities, with some limitations. Microtubule assays often use two colors to distinguish a seed or fiducial marker from free tubulin to determine filament polarity. In its current format, TSOAX and SMAUG analyze a single imaging channel. Further, because the kymographs are generated from segmented filaments, they do not retain information that could identify filament plus or minus ends. Growth or shrinkage can be measured but cannot be reliably assigned to a specific microtubule end without additional information.

The value of faster analysis extends beyond increasing throughput. It allows quantitative measurements to inform the next experiment while a reconstitution assay is still being optimized. For example, the frequency and timing of capping events can reveal whether a protein concentration produces enough events to measure, or whether capping occurs too early or too infrequently for the intended experiment. The same principle applies to other measurements and basic assay conditions, such as slide coating, concentration of surface-linkers, density of crowding reactions, and actin concentrations. Faster analysis makes it possible to identify and adjust conditions before committing to larger or more complex experiments. In this way, quantitative analysis becomes part of assay development rather than a step performed only after an experiment is complete.

More broadly, combining standardized filament detection with automated measurement and targeted user verification provides a practical path toward more scalable and reproducible analysis of actin reconstitution assays. By applying the same segmentation and measurement criteria across filaments, SMAUG reduces the selection and measurement bias inherent to manual analysis. Because the analysis is not tied to a particular assay format, it can be applied across standard open-flow and microfluidics-assisted TIRF-based systems^3,4,49,54^, making quantitative measurements more directly comparable across experiments and laboratories. As reconstitution assays incorporate increasing numbers of regulators and increasingly complex filament behaviors, the ability to analyze large numbers of individual filaments using consistent criteria will become increasingly useful. The modular, open-source design of the TSOAX-SMAUG pairing provides a framework that can be adapted as these experiments become more complex. Rather than replacing user interpretation, this approach decreases the amount of manual selection and measurement required, reduces opportunities for investigator bias, and focuses human effort on the measurements and structures for which biological judgement remains essential.

## Methods

### Reagents

All chemicals and reagents were purchased from Thermo Fisher Scientific (Waltham, MA) unless otherwise indicated.

### Protein expression and purification

All recombinant constructs were sequence-verified by GenScript (Piscataway, NJ). Human profilin-1 (PFN1) in a modified pET vector pMW172^33,55^, mDia1(FH1-C; residues 571-1256) in a modified 6×His-SUMO-containing pET23b vector with flanking AgeI and NotI sequences^21,36^, capping protein (CP) in pET3d^56^, and fascin in pTYB12^46^ were expressed from previously described plasmids. All recombinant proteins were expressed in Rosetta2(DE3) pRare2 cells and induced with 0.4 mM IPTG at 18 ^o^C for 22 h. Untagged proteins (i.e., PFN1 and CP) were purified by anion exchange chromatography followed by size exclusion chromatography on a Superdex75 increase (10/300) (Cytiva, Malborough, MA) in 50 mM (PFN1) or 20 mM (CP) Tris, pH 8, 50 mM KCl, and 1 mM DTT^21,33,35,56,57^. 6×His-SUMO-tagged mDia1(FH1-C) was purified by cobalt affinity chromatography, ULP1 cleavage, and size-exclusion chromatography on a Superose6 (10/300) (Cytiva) in 50 mM HEPES, pH 7.4, 100 mM KCl, 2 mM DTT, 10% glycerol, and 0.02% NP-40^33,36^. Fascin was expressed with an intein-mediated self-cleaving affinity tag, purified by affinity chromatography, and auto-cleaved before size exclusion chromatography in 20 mM Tris pH 8, 100 mM NaCl, and 50 mM DTT^46^. Protein purity was assessed by SDS-PAGE and concentrations determined by Coomassie densitometry relative to BSA standards.

Rabbit skeletal muscle actin was purified from acetone powder or labeled with AlexaFluor 488 NHS ester as described previously^36,58^. Labeling efficiency was determined spectro-scopically at ε_290_, 25,974 M^-1^ cm^-1^ for actin and ε_495_, 71,000 M^-1^ cm^-1^ for AlexaFluor 488, with a correction factor of 0.11. Biotin-actin was purchased from Cytoskeleton, Inc. (Denver, CO) and resuspended in G-buffer according to the manufacturer’s instructions. Labeled actin was stored in 50% glycerol at -20 ^o^C, dialyzed into G-buffer (3 mM Tris, pH 8, 0.5 mM DTT, 0.2 mM ATP, 0.2 mM CaCl_2_) 18 h before use, and pre-cleared by ultracentrifugation at 353,000 × *g* for 1 h before use. Bovine thymus-derived Arp2/3 complex and GST-VCA (N-WASP residues 428-505) were provided by Brad Nolen (University of Oregon)^59^. All proteins were aliquoted, flash frozen, and stored at -80 ^o^C.

### Total Internal Reflection Fluorescence (TIRF) microscopy assays

Flow chambers were assembled from biotin-PEG-coated coverslips (24 × 60 mm, #1.5) adhered to µ-slide VI chambers (IBIDI, Fitchburg, WI) using 0.12 mm SA-S-Secure Seal double-sided tape (GraceBio-labs, Bend, OR) and 5-min Epoxy (Loctite, Rocky Hill, CT)^4^. Chambers were conditioned sequentially with 1% BSA, 0.005 mg/mL streptavidin, 1% BSA, and 1× TIRF buffer (20 mM imidazole [pH 7.4] 50 mM KCl, 1 mM MgCl_2_, 1 mM EGTA, 0.2 mM ATP, 10 mM DTT, 40 mM glucose, 0.25% methylcellulose [4000 cP], 10 mg/mL glucose oxidase, and 2 mg/mL catalase) prior to introduction of actin polymerization reactions^4^. Unless otherwise indicated, reactions contained 1 µM total actin (10% AlexaFluor 488-labeled), including 3.6 nM biotinactin to facilitate surface anchoring. For actin dose-response experiments in Figure 2, total actin concentration was varied as indicated, while biotin-actin concentration and percent label were not. Reactions containing actin binding proteins contained 1 µM actin (10% AlexaFluor 488-labeled) and 1 µM profilin, 5 nM mDia1(FH1-C), 10 nM CP, 100 nM Arp2/3 complex, 500 nM GST-VCA, or 1 µM fascin, in the combinations indicated in the figures and legends. TIRF reactions were initiated by adding actin monomers to the microfuge tube containing all other reaction components,which initiates actin assembly under these conditions.

TIRF images were acquired on a Nikon iLAS TIRF microscope equipped with a 60× 1.49 NA Plan Apo oil-immersion objective using 488 nm excitation (100 ms exposure; 90 mW laser set to 10% power) (Nikon, Inc, Melville, NY) and a Prime BSI express sCMOS camera (Photometrics, Inc, Tucson, AZ), yielding 111.41 µm^2^ field of view (FOV). Time-lapse images were acquired every 5 s for 15 min. Laser power and exposure settings were kept constant within each experiment. Images were denoised using the denoising function in NIS-Elements. These settings were kept constant across conditions within each experiment.

### Computational filament segmentation and quantitative analysis with TSOAX and SMAUG

Filament segmentation and tracking were performed using TSOAX ^9^ followed by analysis with a custom Python-based graphical user interface (GUI), SMAUG. Time-lapse TIRF movies were imported into TSOAX and processed using optimized detection parameters (Supplementary Table 1). TSOAX detects filament centerlines (e.g., “snakes”) in each frame and links them through time to preserve filament identity in an image sequence. TSOAX output files were analyzed using SMAUG, which enables visual overlay of segmented filaments on raw data, user validation of tracking fidelity, and automated extraction of quantitative parameters. Users inspected the computationally tracked filaments overlaid on the raw images and could delete incorrectly tracked filaments or exclude tracking errors. Quantitative measurements were calculated by SMAUG after user validation or when the user declined further filament tracking. For each tracked filament, the software reports length over time (slope), elongation rates from user-defined growth intervals, per-frame filament counts (e.g., nucleation), per-frame filament-filament junction counts (e.g., filament branch points), and mean filament intensity along each segmented filament for intensity-based analyses. SMAUG also generates kymographs of selected actin filaments by extracting the image intensity along the computationally traced filament in each frame and displaying these data as a function of time.

SMAUG can also fit multiple linear segments to filament length versus time trajectories, allowing changes in filament elongation behavior to be identified. The breakpoint locations between these multiple linear segments were determined using the pwlf Python library. The pwlf library determines optimal breakpoint locations by numerically minimizing the sum of squared residuals between the measured filament lengths and the fitted model. The resulting break-points are then also used to fit the corresponding intensity data. SMAUG automatically determines whether a one- and two-segment fit is better using the Bayesian information criterion (BIC), which balances goodness of fit against model complexity. The number of segments and the fitting interval can also be adjusted manually up to 6 sequential segments. We used this approach to quantify single-molecule capping events by capping protein (CP). Filaments that continuously elongated were represented by a single linear fit, whereas filaments containing a change in elongation behavior were represented by multiple linear fits. To be classified as capped, filaments had to exhibit no net growth over 5 consecutive frames. When sorting all filaments tracked by TSOAX in Figures 4J and 4K, filament segments with slopes between - 0.2 and 0.2 pixels per frame were counted as capped filaments if they lasted for a minimum of 3 consecutive frames. The fraction of capped filaments was calculated for each field of view as the number of filaments containing a capped segment divided by the total number of filaments. For visualization and comparison of capping frequency at a defined time point, the percentage of filaments that had undergone capping by 250 s was calculated. For analysis of capping over time, the percentage of filaments that had undergone capping by each time point was calculated and fit to the equation: PFC = PFC_0_ + PFC_max_ - PFC_0_) × (1-e^(-CRC × t)^), where PFC is the percent of capped filaments, PFC_0_ is the percent of capped filaments at the start of sample imaging, PFC_max_ is the maximum percent of capped filaments, and CRC is the filament capping rate constant. Fits were performed using all filaments present in each FOV (n = 9 FOVs, from N = 3 independent reactions ± SD). SMAUG was also used to identify rare uncapping events, which were identified by filaments with at least 3 sequential linear segments corresponding to free elongation, capping, and resumed filament elongation.

SMAUG-based analyses are reported in pixels. SMAUG does not read microscope-specific calibration metadata. Thus, pixel-based measurements require conversion to units based on the microscope setup used for image acquisition. The imaging system described above was used for all datasets associated with this work, and the conversion factor used was 0.1088 µm per pixel. Elongation rates were converted from pixels per frame to µM^-1^ s^-1^ using the pixel calibration and 5 s imaging interval, then to subunits s^-1^ using the established conversion of 370 actin subunits per µm^18^, and normalized to the actin concentration where indicated. Measurements are exported in a tabulated spreadsheet and can be organized by field of view, experimental replicate, or other user-defined organization for downstream analysis and presentation.

### manual verification of software measurements

Measurements were performed from the same TIRF datasets used for semi-automated analysis. Filament nucleation was quantified as the total number of filaments per (111.41 µm)^2^ field of view at 150 s after the initiation of the biochemical reaction. Elongation rate measurements were made in FIJI^60^ by measuring filament length at 6 time points per filament. Slopes of these lines were calculated in Microsoft Excel and converted to subunits s^-1^ µM^-1^ by multiplying by 370^18^. For manual measurements, 17 filaments per field of view (FOV), or all filaments present if fewer, were measured across N = 3 independent reactions, summing to n = 153 filaments per condition. The exception was Supplementary Figure 1B, where 6 filaments per FOV were measured from 3 FOVs per0020condition across N = 3 independent TIRF reactions, yielding n = 54 total filaments per condition. The time required to measure single filament elongation rates was recorded with a stopwatch for each measurement, summed, and averaged across the FOV. For direct comparison, automated datasets were subsampled to match the number of filaments measured manually.

### Statistical analyses

Statistical analyses were performed using GraphPad Prism 11 (version 11.1.0). Statistical tests and replicate numbers are provided in the figure legends. Two-tailed Welch’s t-tests were used for pairwise comparisons and one-way ANOVA with Tukey’s post hoc testing for multiple comparisons. Statistical significance was defined as *α* = 0.05. P-values shown in the figures represent comparisons with the buffer control (actin alone).

## Supporting information

SMAUG_README

SMAUG_GUI_python_code

Table_S1_optimized_parameters

## Data and code availability

The SMAUG GUI, all raw TIRF movies, an optimized TSOAX parameter file, and example datasets for getting started with SMAUG are freely available without restrictions via: 10.5281/zenodo.23224944. The open-source TSOAX segmentation software is available at the Vavylonis laboratory website (https://www.lehigh.edu/~div206/tsoax), and the Python GUI (SMAUG) source code is available on Github (https://github.com/ridillaj/SMAUG_filament_tracking_GUI).

## Supplemental information

The supplement includes 1 Figure and 1 Table.

### Acknowledgements

We acknowledge the use of ChatGPT (OpenAI) for general brainstorming and suggestions regarding approaches to developing the SMAUG graphical user interface. The authors developed, implemented, and validated the resulting code. We are grateful to B. Nolen (University of Oregon) for the Arp2/3 complex and VCA and to B.K. Haarer for thoughtful discussions and suggestions. We thank R.A. Boone for reminding us that every substantial hoard begins with a single piece of gold. This work was supported by the National Institutes of Health GM133485 to J.L. Henty-Ridilla and GM136372 to D. Vavylonis.

## Author contributions

Author Contributions: A.D. Nobles: Conceptualization, Data Curation, Formal Analysis, Investigation, Methodology, Software, Validation, Visualization, Writing: original draft, review, & editing; D.M. Rutkowski: Conceptualization, Methodology, Software, Writing: review & editing; C.R. Belden: Software, Writing: review & editing; L.M. Haney: Investigation, Writing: review & editing; A.E. Davenport: Investigation, Writing: review & editing; D. Vavylonis: Conceptualization, Funding Acquisition, Methodology, Resources, Software, Writing: original, review & editing; J.L. Henty-Ridilla: Conceptualization, Data Curation, Funding Acquisition, Investigation, Methodology, Project Administration, Resources, Supervision, Validation, Visualization, Writing: original, review & editing.

## Supplemental Information

**Supplementary Figure 1.**
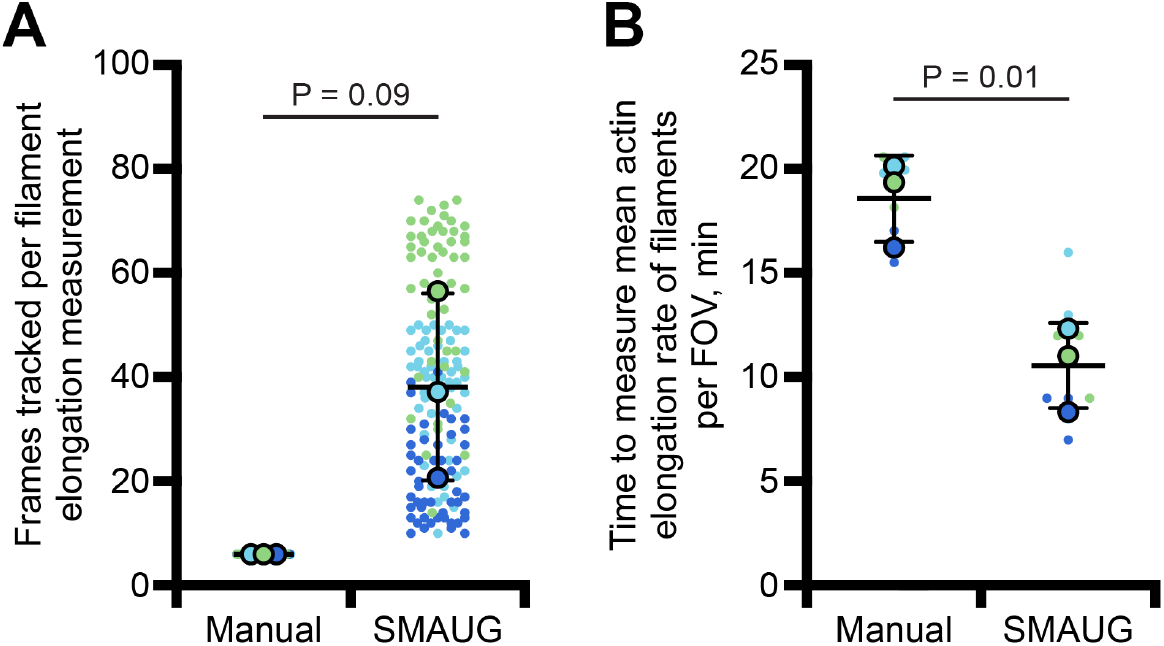
SMAUG enables longer filament tracking and faster measurement of actin elongation rates compared to manual analysis. **(A)** Number of frames tracked per filament for elongation rate measurements performed manually or using SMAUG. **(B)** Time required to measure the mean actin elongation rate for 6 filaments per field of view (FOV) using manual or SMAUG analysis. Measurements were performed for 3 FOVs per condition across N = 3 independent TIRF reactions (n = 54 filaments per condition). Small dots represent individual measurements, and large dots indicate means for each independent reaction. Colors denote independent reactions. Data are mean ± SD. Statistical comparisons between manual and SMAUG measurements were performed using Welch’s t-test (S1A, p = 0.09; S1B, p = 0.01).

**Supplementary Table 1. TSOAX parameters used for analysis of actin filament dynamics.**

Optimized TSOAX parameter settings used to segment and track actin filaments in the experiments shown in each figure. Because the actin filament density and dynamics varied across experimental conditions, TSOAX parameters were optimized for each dataset as indicated.

